# Cytolethal Distending Toxin: from mitotic DNA damage to cGAS-dependent pro-inflammatory response

**DOI:** 10.1101/2020.06.09.141648

**Authors:** Benoît J. Pons, Aurélie Pettes-Duler, Claire Naylies, Frédéric Taieb, Saleha Hashim, Soraya Tadrist, Yannick Lippi, Gladys Mirey, Julien Vignard

## Abstract

The Cytolethal Distending Toxin (CDT) is a bacterial genotoxin that activates the DNA damage response and induces inflammatory signatures in host cells, but the precise relationship between these outcomes has not been addressed so far. CDT induces a singular time-dependent increase of DNA damage and cell cycle defects, questioning on possible impaired response to this toxin over the cell cycle. Here, we identify mitosis as a crucial phase during CDT intoxination. Despite active cell cycle checkpoints and in contrast to other DNA damaging agents, CDT-exposed cells reach mitosis where they accumulate massive DNA damage, resulting in chromosome fragmentation and micronucleus formation. These micronuclei are recognized by cGAS that elicits an inflammatory signature resulting in cell distention and senescence. Our results unravel for the first time the mitotic consequences of CDT genotoxic activity and relate them to pro-inflammatory cellular response. These findings may have important implications during bacterial infection regarding CDT-mediated immunomodulatory and tumorigenic processes.

## Introduction

Several pathogenic bacteria hijack host cell cycle to promote tissue colonization and ensure potent infection by producing specific virulence factors termed as cyclomodulins [1]. These include genotoxins that target chromatin to induce DNA damage-dependent cell cycle defects, among which lies the Cytolethal Distending Toxin (CDT). CDT is produced by more than 30 phylogenetically distant Proteobacteria, such as *Escherichia coli* (*E. col*), *Haemophilus ducreyi* (*H. duc*), *Campylobacter jejuni* (*C. jej*) or *Helicobacter hepaticus* (*H. hep*), responsible for major foodborne diseases all over the world [2]. CDT from *C. jej* or *H. hep* promote persistent gastrointestinal bacterial colonization, accompanied by chronic gastritis and/or typhlocolitis in mice models [3,4]. Besides, the production of CDT from the same bacteria is associated to carcinogenicity in susceptible mice backgrounds [5–7], raising the issue of putative links between CDT-mediated DNA damage, proinflammatory response from the host and tumorigenic processes [8].

CDT adopts an AB_2_ structure [9] where the binding “B_2_” moiety, composed by CdtA and CdtC, mediates delivery of the active “A” subunit, CdtB, into host cell. CdtB belongs to the cation-dependent endonuclease-exonuclease-phosphatase family and has been structurally and functionally related to DNase I, albeit its biochemical activity still needs to be clarified [10]. There is growing evidence that CDT exerts a genotoxic activity in a broad range of host cell lineage [11]. Remarkably, CDT-exposed cells accumulate DNA strand breaks and activate the DNA Damage Response (DDR) through the ATM-CHK2 and ATR-CHK1 axis to promote cell cycle arrest and apoptotic cell death [12–14]. Cells that survive the acute phase of intoxication enter a senescent state, characterized by a permanent cell cycle arrest, persistent DDR activation, enhanced beta-galactosidase (β-Gal) activity and cell distension, the latter having given its name to the toxin [15]. Another general feature of the host cell response to CDT, which is reminiscent to the senescence-associated secretory phenotype, is the production of cytokines [16], notably the pro-inflammatory mediators IL-1β, IL-6 and IL-8, in various cell lineages [15,17,18].

CdtB primarily induces single-strand breaks (SSBs) that degenerate into double-strand breaks (DSBs) at replication forks during S phase, or can directly generate DSBs throughout cell cycle at high CDT doses when two single-strand breaks face each other [19]. Therefore, cells intoxicated with CDT employ a battery of DNA repair mechanisms to maintain cell viability by preventing DSBs accumulation and genetic instability [20]. As CdtB remains active for at least 48 hours once internalized [14,20], the balance between CdtB-induced DNA damage and the repair capacity of the host should dictate the fate of intoxicated cells. In light of this rationale, previous reports pointed out the time-dependent increase of the proportion of cells suffering DSBs and cell cycle arrest during a CDT exposure covering several cell cycles [14,19]. This can be explained by an enhanced CdtB nuclease activity over time, or a decreased DNA repair activity during the course of intoxication. Indeed, DSB repair is specifically inhibited during mitosis through the phosphorylation of RNF8 and 53BP1, preventing their recruitment to damaged chromatin [21]. However, the mitotic consequences of CDT exposure have never been addressed so far.

Here, we explore the mitotic phenotype of human cells exposed to CDT from *E.col* and relate it to micronucleus (MN) induction. We demonstrate that except for highly lethal doses, CDT-exposed cells can enter mitosis without obvious induction of CHK1/CHK2 activation and G2 blockage. Mitotic cells accumulate more rapidly DNA damage than interphasic cells, resulting in mitotic delay at metaphase and MN formation. Contrary to other DNA damaging agents, ATR does not protect CDT-treated cells from such outcomes. The cytosolic DNA sensor cyclic guanosine monophosphate (GMP)-adenosine monophosphate (AMP) synthase cGAS binds to CDT-induced MN, eliciting a pro-inflammatory response and leading to senescence and cell distention. Our data demonstrate for the first time the unique capacity of CDT to overcome the G2 checkpoint and induce DNA damage in mitotic cells, resulting in MN formation to promote cGAS-dependent senescence and pro-inflammatory response.

## Results

### CDT impacts cell viability without obvious DNA Damage Response activation

In an attempt to better characterize its genotoxic potential in human cells, *E.col* CDT has been compared to three DSB-inducing controls, with different mode of action: topoisomerase I and II poisons, respectively campthotecin (campto) and etoposide (etop), and the DNA cross-linking agent mitomycin C (MMC). When HeLa cells are treated with increasing concentrations of the control compounds, a good correlation is observed between clonogenic survival (Fig. 1A) and phosphorylation of H2AX, CHK1 and CHK2 after 24 h (Fig. 1B), as a surrogate of DDR activation. On the other side, 0.025 and 0.25 ng/ml of CDT decrease cell viability despite no obvious phosphorylation of H2AX, CHK1 and CHK2 during the first 24 h, suggesting that, at least for doses lowest than 2.5 ng/ml, CDT-mediated cytotoxicity cannot be attributable to rapid DDR activation and related cell cycle arrest. Whatever the DNA damaging agent, the highest concentration tested induced massive DDR activation and prevented mitotic entry (Fig. S1), resulting in 100% cell death (Fig. 1A). In the same way, previous studies used most of the time CDT concentration around 1 μg/ml and beyond, resulting in substantial cell cycle arrest followed by apoptosis. Currently, the physiological concentration of CDT to which cells are exposed during a natural infection is still unknown [8], but one may expect that the cellular effects caused by moderate concentrations of toxin should be at least as relevant than analyzing highest doses. As we aimed to elucidate the mitotic phenotype of damaged cells and beyond, treatments totally blocking cell cycle were excluded from the rest of the study.

**Figure 1.**
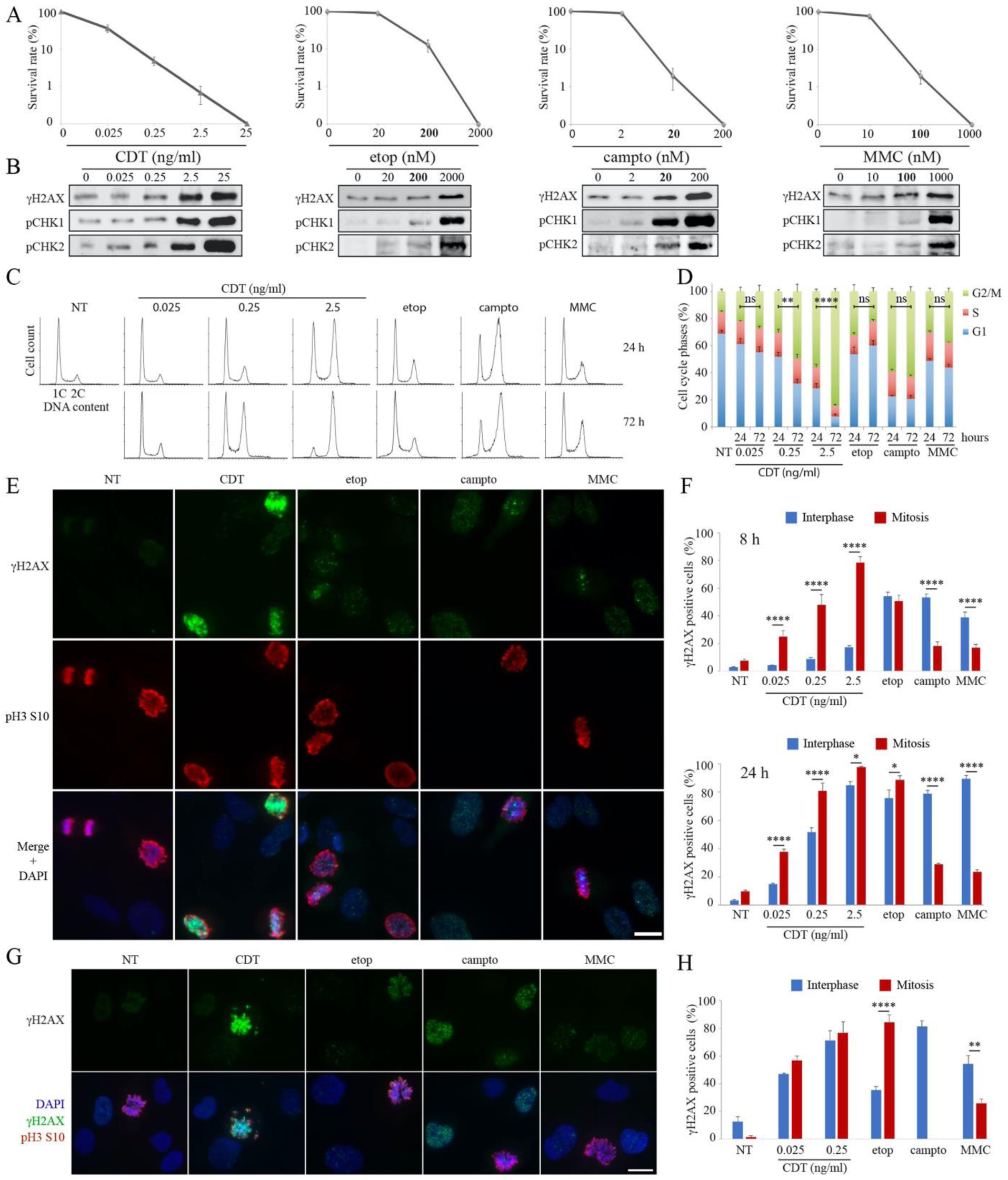
CDT exposure results in early and massive DNA damage in mitotic cells. **(A)** HeLa cells were exposed to CDT, etop, campto or MMC and subjected to colony formation assay. Results present the mean ± SD (*N*≥3). **(B)** HeLa cells were exposed for 24 h to CDT, etop, campto or MMC. Soluble fractions were analyzed by Western blotting. **(C** and **D)** HeLa cells were exposed for 24 or 72 h to CDT, etop 200 nM, campto 20 nM or MMC 100 nM and subjected to cell cycle analyzes by flow-cytometry. Graphs of the cell cycle profiles from one representative experiment (C) and quantitative analyzes of the results (D) are shown (*N*≥3). Statistics were conducted on the G2/M population. (**E** to **H**) HeLa (E and F) or HCEC (G and H) cells were exposed to CDT, etop 200 nM, campto 20 nM or MMC 100 nM for 24 h or for 8 h (HeLa only), and analyzed by immunofluorescence microscopy with antibodies directed against γH2AX and pH3 S10. Representative images (E and G) and quantification (F and H) are shown. Scale bar = 20 μm. (**D, F** and **H**) Data represent the mean ± SEM (*N*≥3). Statistics (only G2/M for D) were calculated by two-way ANOVA followed by Sidak’s multiple comparison test.

### CDT-exposed cells experience DNA damage at mitosis

Similar to DNA damage, CDT-induced cell cycle arrest significantly increases from 24 to 72 h, as previously reported [19], whereas cells exposed to etop 200 nM, campto 20 nM or MMC 100 nM exhibit stable cell cycle defects over time (Fig. 1, C and D). These results imply that at least a part of CDT-exposed cells blocks their cell cycle after the first round of cell division, suggesting they can initially progress through mitosis before induction of DNA damage. Strikingly, asynchronous cells exposed to CDT display an intense γH2AX signal in mitosis compared to interphase or to the few γH2AX foci observed in mitotic cells exposed to the other DNA damaging agents (Fig. 1E). The basal DNA damage-independent γH2AX signal present in unchallenged mitotic cells [22], diffuse all along the condensed chromosomes from prometaphase to anaphase, is easily distinguishable from the signal induced by CDT or the genotoxic controls. Moreover, mitotic cells represent the first population to be damaged during the course of CDT treatment, contrary to campto or MMC (Fig. 1F). Concerning etop, mitotic cells are equally affected after 8 h compared to interphasic cells, but only display discrete γH2AX foci. Etop at highest concentration of 10 μM has previously been shown to induce important DNA damage on mitotic HeLa cells, but a synchronization in mitosis is required to ensure that an important proportion of cells will be affected [23].

The huge γH2AX increase in CDT-exposed mitotic cells represents a general cellular response and is observed with CDT from other bacterial origins or with other cancer cell lines (Fig. S2). Moreover, normal human colonic epithelial cells (HCECs) that we have previously shown to be susceptible to CDT intoxination [24], also exhibit a similar response (Fig. 1, G and H). Thus, our results reveal that mitotic cells are particularly sensitive to CDT genotoxicity, probably because DSB repair is inhibited during mitosis through the phosphorylation of several DDR factors by CDK1 and PLK1 kinases [21].

### CDT induces DNA double-strand breaks during mitosis

The strong γH2AX increase after CDT can be observed all along the mitotic phases (Fig. 2, A and B). In addition, a dose dependent increase of chromosome fragments that does not properly align during metaphase or segregate at anaphase can be observed. To exclude that this γH2AX signal results from DNA damage induced before mitotic entry, HeLa cells were enriched in mitosis by a 16 h nocodazole block before to be co-exposed for 6 h to nocodazole and CDT or genotoxic control agents. Similar to asynchronous cells, cells treated with CDT during mitosis exhibit a strong γH2AX level compared to etop, campto or MMC (Fig. 2C). This CDT-induced γH2AX signal depends both on ATM and ATR, as stated by the use of specific inhibitors, respectively KU-55933 (ATMi) and VE-821 (ATRi) (Fig. 2D). Finally, to confirm that the γH2AX mitotic signal depends on DSB induction, cells arrested in mitosis were exposed to CDT or control genotoxic compounds before to be subjected to neutral comet assay (Fig. 2, E and F). Only mitotic cells treated with 0.25 or 2.5 ng/ml of CDT show a significant increase in comet tail moment. As noted above, substantial DNA fragmentation during mitosis has already been observed in response to etop, but at concentrations that normally induce G2 arrest and therefore prevent mitotic entry [23]. Taken together, these data demonstrate that CDT can induce DSB during mitosis that are signaled by ATM and ATR and lead to chromosome fragmentation and missegregation.

**Figure 2.**
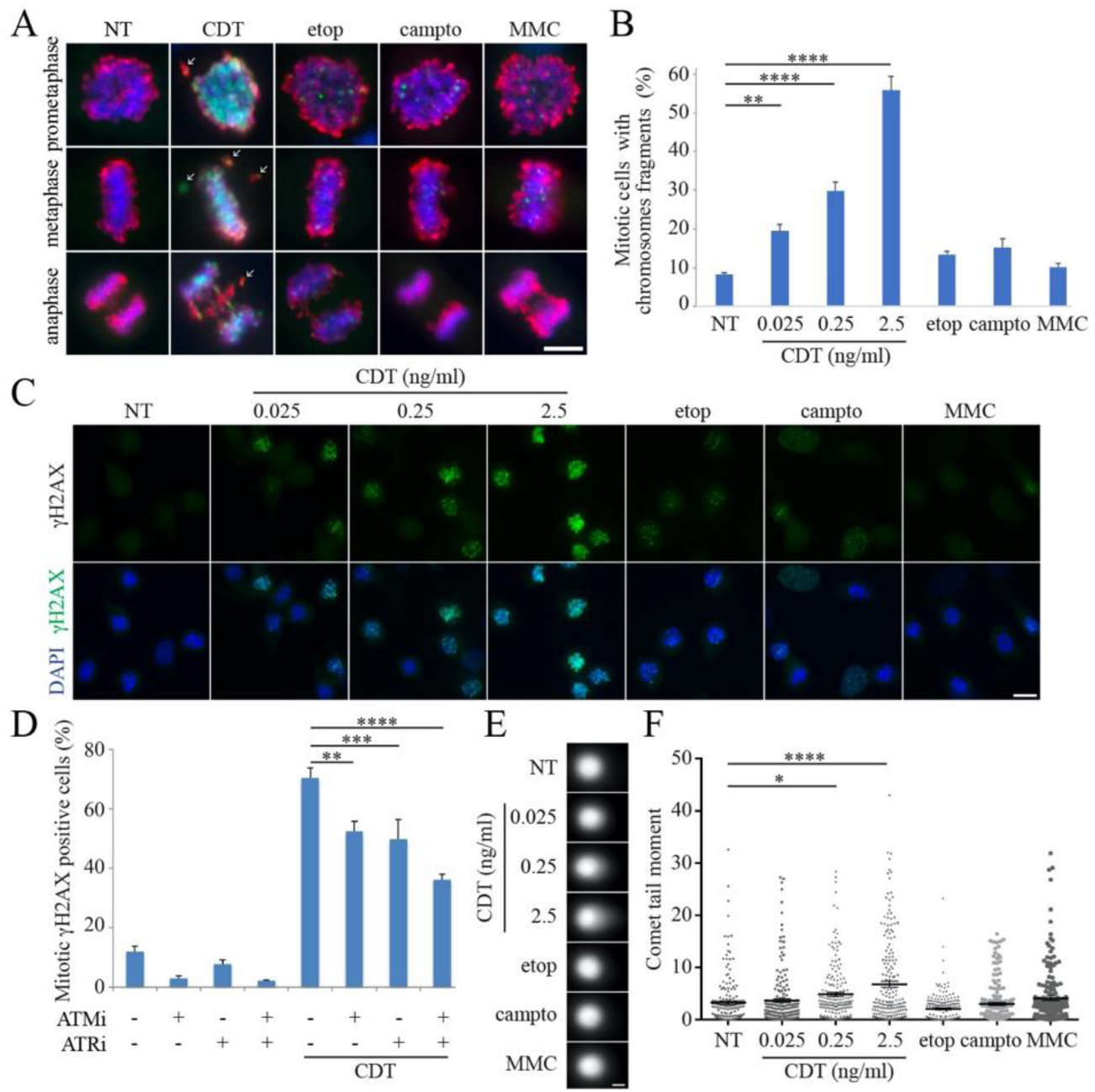
CDT induces DNA damage during mitosis. (**A** and **B**) HeLa cells were exposed for 24 h to CDT, etop 200 nM, campto 20 nM or MMC 100 nM and analyzed by immunofluorescence microscopy with antibodies directed against γH2AX (green) and pH3 S10 (red). Representative images of mitotic cells in prometaphase, metaphase or anaphase (A) and quantification of chromosome fragments in prometaphase and metaphase (B) are shown. Scale bar = 10 μm. White arrows indicate chromosome fragments. (**C**) HeLa cells were exposed for 22 h to nocodazole, with CDT, etop 200 nM, campto 20 nM or MMC 100 nM during the last 6 h and analyzed by immunofluorescence microscopy with an antibody directed against γH2AX. Images of one representative experiment are shown (N=4). Scale bar = 20 μm. (**D**) HeLa cells were exposed for 22 h to nocodazole, with CDT 2.5 ng/ml, ATMi and/or ATRi during the last 6 h and analyzed by immunofluorescence microscopy with an antibody directed against γH2AX. (**E** and **F**) HeLa cells were synchronized in mitosis, exposed for 12 h to CDT, etop 200 nM, campto 20 nM or MMC 100 nM and analyzed by neutral comet assay. Representative images (E) and quantification of individual cells tail moment (F) are shown. Scale bar = 20 μm. (**B, D** and **F**) Data represent the mean ± SEM (*N*=3). Statistics were calculated by one-way (B and F) or two-way (D) ANOVA followed by Dunnett’s multiple comparison test.

### CDT-exposed cells accumulate at mitosis despite active G2 checkpoint

We next monitored more deeply the cell cycle consequences of the CDT-induced DNA lesions. Cell cycle analyses were conducted after immunostaining with antibodies directed against pH3 S10, to identify mitotic cells, and γH2AX. After a 24 h exposure to etop 200 nM, campto 20 nM or MMC 100 nM, cells that progress to mitosis are devoid of γH2AX staining (Fig. 3A), whereas G2 arrest is more pronounced at 10-fold higher concentrations, preventing mitotic entry in agreement with the principle that DDR ensures cells do not enter mitosis with damaged DNA. Indeed, while mitosis coordinates the proper segregation of sister chromatids, controlling chromosome integrity prior separation is performed at the previous G2 phase and implies cell cycle arrest and DNA repair to impede transition to mitosis with damaged DNA [25]. As duration of M phase is short and constant [26], only a very minor part of cells from an asynchronous population may suffer DNA damage at mitosis. However, cells treated with CDT present a dose-dependent augmentation of the population co-stained with pH3 S10 and γH2AX antibodies, representing a 12-fold increase at 2.5 ng/ml of CDT compared to control cells (Fig. 3A). These data strengthen the assumption that CDT genotoxic activity does not preclude mitotic entry due to G2 checkpoint activation.

**Figure 3.**
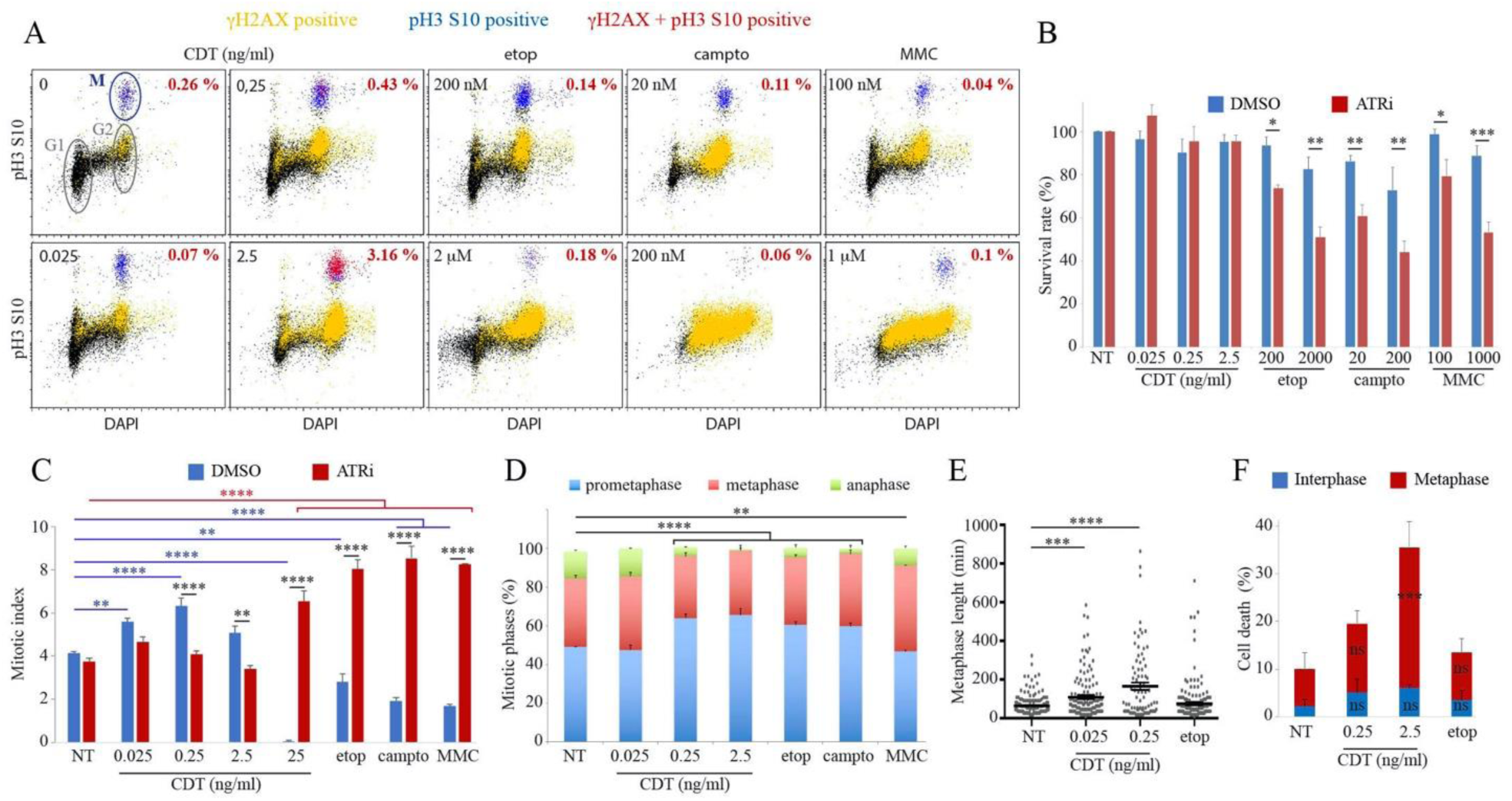
Mitotic phenotype of CDT-exposed cells. (**A**) HeLa cells were exposed for 24 h to CDT, etop, campto or MMC, immunostained with antibodies directed against γH2AX and pH3 S10 and subjected to cell cycle analyzes by flow-cytometry. Graphs of the cell cycle profiles from one representative experiment (*N*=2) show unstained cells (black), or with γH2AX (yellow), pH3 S10 (M for mitosis, blue) or both staining (red). Percentage in red indicate the proportion of γH2AX and pH3 S10 positive cells. (**B**) HeLa cells were exposed for 24 h to CDT, etop, campto or MMC with or without ATRi and subjected to CellTiter-Glo viability assay. Data represent the mean ± SEM (*N*≥3). Statistics were calculated by two-way ANOVA followed by Fisher’s multiple comparison test. (**C**) HeLa cells were exposed for 24 h to CDT, etop 200 nM, campto 20 nM or MMC 100 nM with or without ATRi and analyzed by immunofluorescence microscopy with an antibody directed against pH3 S10 for mitotic index quantification. Data represent the mean ± SEM (*N*≥3). Statistics were calculated by two-way ANOVA followed by Sidak’s multiple comparison test. (**D**) HeLa cells were treated as in C without ATRi and the proportion of prometaphase, metaphase or anaphase were quantified. Statistics (anaphase only) were calculated by one-way ANOVA followed by Dunnett’s multiple comparison test. (**E** and **F**) HeLa cells stably expressing the chromatibody-GFP [30] were exposed to CDT or etop 200 nM and analyzed by time-laps fluorescence imaging. Graphs show the average time duration of metaphase to anaphase transition ± SEM of one representative experiment (N=3) (E), and the mean of cell death at interphase or metaphase ± SEM (*N*=3) (F). Statistics were calculated by one-way (E) or two-way (F) ANOVA followed by Dunnett’s multiple comparison test.

To test this hypothesis, cells were co-exposed with ATRi, given that under these conditions, the G2/M checkpoint induced by CDT or the three DNA damaging controls mostly depends on ATR rather than ATM (Fig. S3A), as noticed for γH2AX in interphasic cells except with etop (Fig. S3B). No viability defect could be observed after only 24 h of CDT, even in the presence of ATRi, while ATR inactivation sensitizes to etop, campto or MMC (Fig. 3B). Moreover, contrary to the mitotic index drop expected in response to a genotoxic insult that induces the G2 checkpoint observable with the control treatments, the mitotic population significantly increases after 0.025 and 0.25 ng/ml of CDT, before slightly decreasing at 2.5 ng/ml (Fig. 3C). Increasing CDT concentration to 25 ng/ml completely prevents mitotic entry, as observed for the control agents and in accordance with DDR activation (Fig. 1B and Fig. S1B). Of note, mitotic index of CDT-exposed HCEC does not significantly decreases, contrary to the other tested genotoxic compounds (Fig. S4A), emphasizing that this specific response to CDT is not limited to HeLa cells. Thus, until a certain threshold is reached, CDT-mediated DNA damage does not firmly activate the G2 checkpoint but rather induces a mitotic delay. Indeed, ATR inhibition only alleviates G2 block with control treatments or with CDT 25 ng/ml, enabling cells to accumulate in mitosis with damaged DNA (Fig. S5), whereas ATRi reduces the mitotic index at lowest CDT concentrations. In summary, our data demonstrate that the ATR-dependent G2 checkpoint does not prevent mitotic entry of cells exposed to moderated CDT concentrations, enabling CDT to induce DNA damage and a lag in mitosis. This atypical scenario is probably a consequence of the particular CdtB genotoxic potential. Indeed, contrary to irradiations that instantaneously induce massive DNA damage, or to chemicals and metabolites from pathogens with limited stability or capacity to induce repeated lesions, CdtB may exert unceasing genotoxic attacks, given that this nuclease remains active for at least 48 h after cellular internalization [14,20]. CDT is the only characterized encoded toxin from mammalian pathogens that has evolved to create DSBs in host genomic DNA [27]. This continuous nuclease activity is opposed to the host DNA repair machinery, implying that under a certain threshold, CDT-induced DNA damage does not obviously activate the DDR and therefore allows G2-M transition (Fig. 1, A-C). Besides, it is noteworthy that other genotoxic compounds finally induce a similar mitotic phenotype that is delayed in time. Indeed, HT-29 cells arrested in G2 after a 25 nM treatment of campto, which is comparable to the present study, finally reach mitosis from 48 h post-treatment with highly damaged chromatin [28]. Such recovery from G2 arrest in the presence of DNA damage has been observed with many DNA damaging agents and is referred to as checkpoint adaptation [29]. Therefore, CDT exposure globally recapitulates the phenotype of cells subjected to checkpoint adaptation, but with more rapid kinetics due to an absence of ATR-dependent G2 checkpoint activation.

### CDT induces metaphase delay and cell death

The mitotic delay observed in response to CDT is characterized by a dose-dependent diminution of the anaphase population, a phenomenon also observed in HCEC (Fig. S6A) or in cells that reach mitosis despite exposure to the genotoxic controls (Fig. 3D). In order to gain insight into the mitotic phenotype of CDT-treated cells, live-cell imaging has been performed on HeLa cells stably expressing the chromatibody fused to GFP, enabling chromatin visualization in real-time [30]. When measuring the timing needed to complete metaphase, we found that unperturbed mitosis takes an average of 64 min that significantly increases to 109 min with 0.25 ng/ml and 164 min with 2.5 ng/ml of CDT, a delay that could not be observed with etop at the moderate concentration of 200 nm (Fig.3E). This phenotype has also been observed with other genotoxic compounds in cells suffering extensive DNA damage [31] and is similar to cells depleted for the mitotic resolvase GEN1 [32,33]. Indeed, mitotic duration is governed by the capacity of cells to dissolve or resolve Hollyday junctions [34], questioning on putative targeting by CdtB of homologous recombination intermediates formed after CDT-induced replicative stress [20], which are processed later in mitosis. Then, monitoring cell death during the course of live imaging revealed that an important part of CDT-exposed cells preferentially dies during metaphase (Fig. 3F). In conclusion, we demonstrate that mitotic cells are particularly affected during CDT intoxination, as evidenced by a prolonged metaphase duration that may eventually result in cell death.

### cGAS recognizes CDT-mediated micronuclei to elicit senescence and inflammatory signature

One of the most well-established consequence of DNA damage is the presence of MN in the next cell generation, resulting from missegregated chromosome fragments during mitosis [29]. CDT induces an ATR-independent MN increase at 0.025 and 0.25 ng/ml (Fig. 4A) that correlates with mitotic index (Fig. 3C). MN induction falls down while increasing CDT concentration as the G2 checkpoint is progressively activated. Indeed, ATR inhibition relieves the G2 block to allow massive MN formation at CDT concentration from 2.5 ng/ml or with DNA damaging controls. CDT-mediated MN increase is also observed in normal HCEC (Fig. S6B). Thus, CDT exposure leads to MN generation despite the presence of active cell cycle checkpoints. To explore the long-term consequence of CDT intoxination, HeLa cells were chronically exposed to 0.25 ng/ml of CDT, inducing more than 95% cell death (Fig. 1A). The surviving fraction was cultured and individual clones were selected as well as a pool of resistant cells (Fig. 4B). The chronically exposed cells were unresponsive to CDT-mediated G2 checkpoint (Fig. 4C), suggesting an adaptation to the toxin, and exhibited an important fraction of micronucleated cells (Fig. 4 D). These cells were subjected to transcriptomic analyses and compared to two control groups, *i.e.* cells without treatment or chronically exposed to a CDT catalytic dead mutant that cannot activate the DDR [19,20]. As depicted in the heatmap resuming expression profile of 9703 significantly regulated genes between the 3 conditions, individual clones and the pool of cells chronically exposed to active wild-type (WT) CDT share a common transcriptional adaptation, whereas the two control groups (non-treated and treated with mutant CDT) cannot be distinguished (Fig. 4E). The majority of the most upregulated genes, when comparing the three groups, depends on the catalytic activity of CDT rather than the presence of the toxin solely (Fig. 4.F). The most upregulated biological processes in cells chronically exposed to WT CDT mainly rely on pro-inflammatory responses, more particularly to type I interferon (IFN) signaling (Fig. 4G). Recent reports have demonstrated that MN recognition by cGAS triggers innate immune activation related to type I IFN signature [35,36]. Interestingly, the proportion of cGAS positive MN increases significantly after 72 h of CDT, compared to the other genotoxic treatments (Fig. 4, H and I). Since MN induction is delayed due to checkpoint adaptation after exposure to etop, campto or MMC, this should explain why cGAS is only recruited to CDT-mediated MN at 72 h. Indeed, cGAS can access to genomic DNA contained in MN only after rupture of their nuclear envelope [36], suggesting the envelope is still intact at 72 h in MN formed after checkpoint adaptation, whereas those derived from CDT intoxination form earlier and exhibit collapsed envelope at the timing of observation. In a comparable situation, irradiated MCF10A cells escape from the G2 checkpoint after 3 days, resulting in MN formation that elicits a cGAS-dependent inflammatory signature after 6 days [35].

**Fig. 4.**
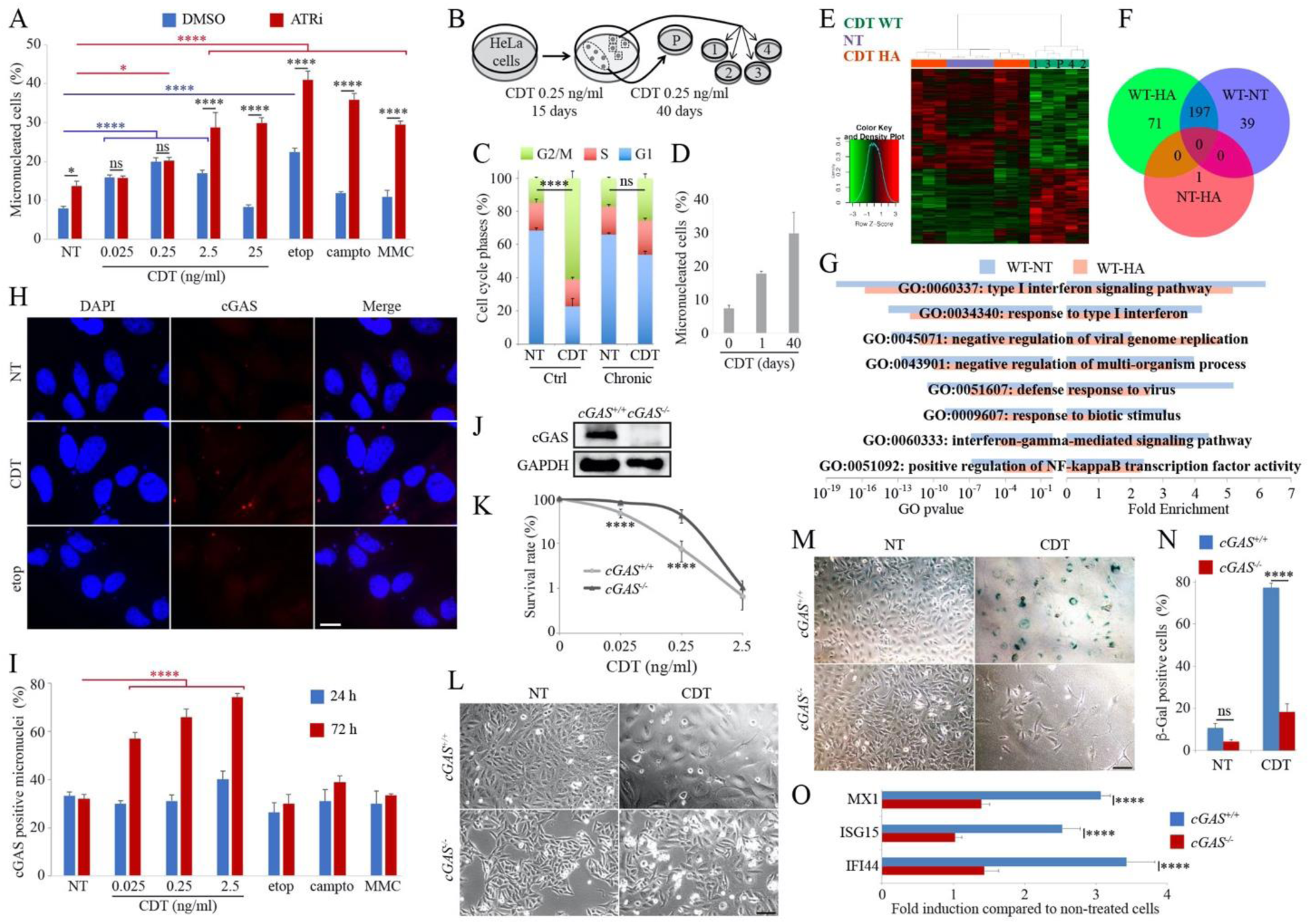
CDT-induced micronuclei promote a cGAS-dependent proinflammatory response. (**A**) HeLa cells were exposed for 24 h to CDT, etop 200 nM, campto 20 nM or MMC 100 nM with or without ATRi and the frequency of cells with micronuclei was quantified. (**B**) Schematic of the selection procedure of HeLa cells chronically exposed to CDT. Individual clones (1-4) and a multiclonal cell population (P) were obtained. (**C**) Cells from B were exposed to CDT 2.5 ng/ml and subjected to cell cycle analyzes by flow-cytometry. Statistics were conducted on the G2/M population. (**D**) HeLa cells were exposed for 24 h or 40 days as in B and the frequency of cells with micronuclei was quantified. Data represent the mean ± SEM (*N*≥3). (**E-G**) Cells non-treated (NT) or from B exposed to wild-type CDT (CDT WT) or the catalytic mutant H153A (CDT HA) were compared after microarray gene expression analysis. Heatmap (E) shows the significantly altered genes expression (FDR<0.05 and fold change>1), and Venn diagram (F) shows overlap between the most upregulated genes (FDR<0.05 and fold change>3). Graphs (G) represent the 8 most significant upregulated biological processes sorted according to adjusted *p* value obtained from a hypergeometric enrichment test on Gene Ontology (GO) processes. (**H** and **I**) HeLa cells were exposed for 24 or 72 h to CDT, etop 200 nM, campto 20 nM or MMC 100 nM and analyzed by immunofluorescence microscopy with an antibody directed against cGAS (red). Representative images at 72 h (H) and quantification of cGAS positive MN (I) are shown. Scale bar = 20 μm. Data represent the mean ± SEM (*N*≥3). Statistics were calculated by two-way ANOVA followed by Dunnett’s multiple comparison test. (**J**) Soluble fractions of *cGAS*^*+/+*^ and *cGAS*^*-/-*^ HeLa cells were analyzed by Western blotting. (**K**) *cGAS*^*+/+*^ and *cGAS*^*-/-*^ HeLa cells were exposed to CDT and subjected to colony formation assay. (**L**-**N**) *cGAS*^*+/+*^ and *cGAS*^*-/-*^ HeLa cells were exposed to CDT 2.5 ng/ml for 90 h and analyzed by light microscopy without (L) or with β-Gal staining (M and N). Scale bar = 10 μm. Quantification of β-Gal positive cells is shown (N). (**O**) *cGAS*^*+/+*^ and *cGAS*^*-/-*^ HeLa cells were exposed to CDT 0.25 ng/ml for 10 days and the mRNA level of *MX1, ISG15* and *IFI44* was analyzed by RT-qPCR. (**A, C, K, N** and **O**) Data represent the mean ± SEM (*N*≥3). (**A, C, K, N** and **O**) Data represent the mean ± SEM of at least three independent experiments. Statistics (only G2/M for C) were calculated by two-way ANOVA followed by Sidak’s multiple comparison test.

The cGAS-STING axis activates IRF3 and NF-κB, two transcriptional inducers of type I IFNs and other cytokines [37]. cGAS has also been shown to play a crucial role during cellular senescence induced by DNA damage [38]. To better understand the role of cGAS in response to CDT injury, cGAS knockout HeLa cells (*cGAS*^*-/-*^) were generated (Fig. 4J). *cGAS*^*-/-*^ cells are more resistant to low CDT concentrations (0.025 and 0.25 ng/ml) than their WT cGAS counterpart (Fig. 4K). Moreover, despite similar viability loss between *cGAS*^*+/+*^ and *cGAS*^*-/-*^ cells at 2.5 ng/ml of CDT, a 72 h exposure does not induce cell distention (Fig. 4L) nor β-Gal staining as a marker of senescence in cGAS-depleted cells (Fig. 4, M and N). Finally, the basal and CDT-induced mRNA expression of three type I IFN signaling target genes (*MX1, ISG15* and *IFI44*) was compared between *cGAS*^*+/+*^ and *cGAS*^*-/-*^ cells after 10 days. An approximately 3-fold increased expression level of each tested gene after chronic CDT exposure was observed, but absent in *cGAS*^*-/-*^ cells (Fig. 4O). Altogether, these results unravel the essential role of cGAS during CDT intoxination through MN recognition, eliciting cell distention, senescence and pro-inflammatory responses. However, we cannot exclude that in CDT-exposed cells, cGAS inhibits DSB repair and/or promotes mitotic cell death upon retardation of metaphase-anaphase transition [39,40], therefore participating in the sensitivity to the toxin.

Persistent infection causes chronic inflammation, a driving force of tumor development [41]. CDT production during bacterial infection promotes inflammation and persistent colonization, and may eventually contribute to tumorigenesis [3–7]. In a broad range of cellular lineages, CDT intoxination induces DNA damage and a proinflammatory signature [8], but how these two events are linked has never been addressed so far. Our findings identify cGAS as the missing piece between CDT genotoxic and immunomodulatory activities. cGAS is a key modulator of the innate immune response that can influence the tumor microenvironment in ways that may be detrimental or beneficial [42]. Further investigations will be necessary to unravel the role of the cGAS-dependent surveillance of micronuclei induced during natural infection with CDT producing bacteria.

## Materials and Methods

### Cell culture and treatments

HeLa, U2OS and HCT116 cells were cultured in DMEM (Life Technologies), supplemented with 10% heat-inactivated fetal bovine serum (FBS, Gibco) and 1% antibiotics (penicillin/streptomycin). Caco-2 cells were cultured in DMEM supplemented with 20% heat-inactivated FBS and 1% antibiotics. The HepaRG cells (Biopredic, France) were cultured in William’s E medium (Life Technologies) supplemented with 10% of FBS, 1% antibiotics, 5 μg/ml insulin, 2mM l-glutamine and 50 μM hydrocortisone hemisuccinate. Human colonic epithelial cell (HCECs), generated and provided by Pr Jerry W Shay, were cultured as previously described [24] in 4:1 high-glucose DMEM/medium 199 supplemented with 2% FBS, epidermal growth factor (20 ng/ml), hydrocortisone (1 mg/ml), insulin (10 mg/ml), transferrin (2 mg/ml), sodium selenite (5 nM), and Gentamycin sulfate (50 μg/ml). Cells were maintained at 37°C in a humidified atmosphere containing 5% CO_2_, and subcultured approximately every 2–3 days.

*E.col* CDT (WT or the catalytic dead mutant H153A), *H. duc* CDT and *C. jej* CDT were produced and purified as previously described [7,10,19]. ATM inhibitor KU-55933 (ATMi) and ATR inhibitor VE-821 (ATRi) were purchased from Selleckchem and used at a final concentration of 10 mM. All other chemicals and reagents were purchased from Sigma -Aldrich.

### Western blot analyses

Cells were incubated on ice for 30 min in lysis buffer (50 mM Tris-HCl pH7.5, 500 mM NaCl and 0.5% NP40) containing the HaltTM Protease & Phosphatase inhibitor cocktail (Thermo Scientific) and sonicated on a VibraCell 72434 (Bioblock Scientific). Cell lysates were centrifugated and the supernatant containing total soluble proteins was kept. Proteins were separated by SDS-PAGE and transferred to a nitrocellulose membrane (Amersham). Membranes were incubated with the primary antibody over-night at 4°C. γH2AX antibody was purchased from Merck/Millipore (05– 636), pChk1 (133D3), pChk2 (C13C1), pH3 S10 (D2C8) and cGAS (D1D3G) antibodies from Cell Signaling and GAPDH (GTX100118) antibody from GeneTex. The secondary anti-mouse or anti-rabbit HRP-conjugated antibodies (Jackson Immunoresearch laboratories) were incubated for 1 h at room temperature. Proteins were visualized with the enhanced chemiluminescence substrate ECL (Biorad) and imaged using the ChemiDoc XRS Biorad Imager and Image Lab Software.

### Immunofluorescence analyses and micronucleus assay

Cells were grown on glass coverslips. After at least 24 h of culture, cells were fixed with 4% paraformaldehyde, permeabilized with 0.5% Triton X-100, blocked with 3% BSA and 0.05% IGEPAL, and stained with primary antibodies for 2 h at room temperature in blocking solution (all solutions were prepared in PBS). Cells were washed three times with PBS 0.05% IGEPAL and incubated with the secondary antibodies for 1 h (Alexa Fluor 488 Goat anti-mouse (A32723) or Alexa Fluor 594 anti-rabbit (A32740), purchased from Invitrogen). DNA was stained with 4.6-diamino-2-phenyl indole (DAPI) 30 nM. Coverslips were mounted onto slides with PBS-glycerol (90%) containing 1 mg/ml paraphenylenediamine and observed at 40x magnification with a Nikon 50i fluorescence microscope equipped with a Luca S camera. Cells were counted positive for foci formation when > 10 foci/interphase nuclei or when > 5 foci/mitotic nuclei were detected.

### Cell viability assays

For clonogenic assay, cells were plated in triplicate at a density of 300 to 3000 cells per well in 6 wells plate. One day after seeding, cells were treated and grown for 10 days. Formed colonies were fixed and stained with a 0.25% methylene blue and 100% methanol solution. Colonies containing more than 50 cells were counted and the surviving rate calculated.

For rapid viability testing at 24 h, cells were plated in triplicate at a density of 10000 per well in 96 wells plate. One day after seeding, cells were treated and grown for 24 h. Viability was assessed using the CellTiter-Glo® Luminescent Cell Viability Assay (Promega) according to the manufacturer’s instructions.

### Senescence assay

The senescence-associated β-galactosidase staining was performed using the Senescence β-Galactosidase Staining Kit (Cell Signaling) according to the manufacturer’s instructions. Images were captured by light microscopy.

### Flow cytometry analyses

Cells were fixed with 4% paraformaldehyde for 15 min at room temperature in PBS. For immunostaining, fixed cells were permeabilized with 0.2% triton X-100 and 1% BSA in PBS for 20 min at room temperature. Antibodies against γH2AX and pH3 S10 were incubated for 2 h in 1% BSA at room temperature. After three washes, cells were incubated with the secondary antibodies Alexa Fluor 488 anti-mouse and Alexa Fluor 594 anti-rabbit for 30 min at room temperature. Then, cells were washed three times and incubated in PBS containing DAPI (1 μg/mL) for 15 min before samples were processed using flow cytometry (MACSQuant, Miltenyi Biotec). At least 10000 events were analyzed per sample using FlowLogic software (Miltenyi Biotec).

### Comet assay on mitotic cells

Cells were treated for 18 h with 2 mM thymidine, washed twice with PBS and released in fresh media for 9 h. Cells were treated again with 2 mM thymidine for 17h to be synchronized in early S, washed twice with PBS and released in fresh media for 6 h before adding 50 ng/ml nocodazole. After 4 h, mitotic cells were collected by shake-off and exposed for 12 h to CDT or the genotoxic controls in presence of nocodazole. Comet assay was performed in neutral conditions using the Comet SCGE assay kit (Enzo) according to the manufacturer’s instructions. At least 60 cells were analyzed per sample using OpenComet software.

### Time-lapse imaging

HeLa cells stably expressing the chromatibody-GFP to visualize chromatin in living cells [30] were generated by TALEN insertion at AAVS1 site with co-transfection of SHDP-CMV-VHH-HA-GFP donor plasmid and AAVS1 right and left Talen plasmids (genome TALER AAVS1 safe harbor cloning kit, Genecopoeia) using TransIT-LT1 (MirusBio) according to manufacturer’s instructions. Cells were plated in 4-compartments Petri dish (CellView Greiner Bio-One). Immediately after treatment, time-lapse fluorescent microscopy was performed with a Zeiss Axio-observer inverted videomicroscope with controlled temperature (37 °C), CO2 level (5 %) and humidity. Images were acquired for 90 h every 9 minutes with an AxioCam 506 camera (Zeiss) in two focal planes 5µM apart from each other. Illumination has been set to 75 ms with 20 % laser intensity to reduce phototoxicity. The two focal planes were flattened and 100 to 200 mitosis per sample were analyzed with FIJI software from 4 to 90 h post-exposure to measure the duration of mitotic phases and cell death.

### Generation of cGAS knock-out HeLa cells

The CRISPR plasmid was a generous gift from J. Dupuy (Toxalim, France). The donor plasmid (pCMV-Cas9-GFP, Sigma), containing the sgRNA, Cas9 and GFP, was modified to substitute the sgRNA sequence for cGAS sgRNA 1 (5’-GCTTCCGCACGGAATGCCAGG) or cGAS sgRNA 2 (5’-CGATGGATCCCACCGAGTCT) targeting the first exon of the *cGAS* gene, using the Q5 site-directed mutagenesis kit (New England Biolabs) according to manufacturer’s instructions. The two plasmids bearing cGAS sgRNA1 and 2 were co-transfected in HeLa cells using TransIT-LT1 (MirusBio) according to manufacturer’s instructions. Individual clones were selected by cGAS immunofluorescence and confirmed by Western blotting. Two confirmed *cGAS*^*-/-*^ clones were used in this study.

### Gene expression

Total RNA was extracted with TRIzol reagent (Invitrogen, Carlsbad, CA, USA) mRNA were extracted and the quality of these samples was assessed (Agilent RNA 6000 Nano Kit Quick, Agilent Bioanalyzer 2100); RNA Integrity Number (RIN) of these mRNA was superior to 9.8.

For transcriptomic data, gene expression profiles were obtained at the GeT-TRiX facility (GénoToul, Génopole Toulouse Midi-Pyrénées, France) using Agilent SurePrint G3 Human GE v2 microarrays (8×60K, design 039494) following the manufacturer’s instructions. For each sample, Cyanine-3 (Cy3) labeled cRNA was prepared from 200 ng of total RNA using the One-Color Quick Amp Labeling kit (Agilent Technologies) according to the manufacturer’s instructions, followed by Agencourt RNAClean XP (Agencourt Bioscience Corporation, Beverly, Massachusetts). Dye incorporation and cRNA yield were checked using Dropsense™ 96 UV/VIS droplet reader (Trinean, Belgium). 600 ng of Cy3-labelled cRNA were hybridized on the microarray slides following the manufacturer’s instructions. Immediately after washing, the slides were scanned on Agilent G2505C Microarray Scanner using Agilent Scan Control A.8.5.1 software and fluorescence signal extracted using Agilent Feature Extraction software v10.10.1.1 with default parameters.

For real-time quantitative polymerase chain reaction (qPCR), 2 μg RNA samples were reverse-transcribed using the High-Capacity cDNA Reverse Transcription Kit (Applied Biosystems). qPCR was performed using the Power SYBR® Green PCR Master Mix and an ABI Prism 7300 Sequence Detection System instrument and software (Applied Biosystems). The following gene-specific primers were used: MX1 (forward: 5’-TTCAGCACCTGATGGCCTA; reverse: 5’-AAAGGGATGTGGCTGGAGAT), ISG15 (forward: 5’-GCGAACTCATCTTTGCCAGTA; reverse: 5’-CCAGCATCTTCACCGTCAG), IFI44 (forward: 5’-ATGGCAGTGACAACTCGTTTG; reverse: 5’-TCCTGGTAACTCTCTTCTGCATA) and TBP1 (forward: 5′-TGTATCCACAGTGAATCTTGGTTG; reverse: 5′-GGTTCGTGGCTCTCTTATCCTC). All samples were run in triplicate. qPCR data were normalized to TBP1 mRNA levels and analyzed with LinRegPCR.v2015.3.

### Statistical analysis

Microarray data were analyzed using R and Bioconductor packages [43]. Raw data (median signal intensity) were filtered, log2 transformed, and normalized using quantile method [44]. A model was fitted using the limma lmFit function [45]. Pair-wise comparisons between biological conditions were applied using specific contrasts. A correction for multiple testing was applied using Benjamini-Hochberg procedure to control the False Discovery Rate (FDR). Probes with FDR≤0.05 were considered to be differentially expressed between conditions. Hierarchical clustering was applied to the samples and the differentially expressed probes using 1-Pearson correlation coefficient as distance and Ward’s criterion for agglomeration.

Other statistical analyses were assessed using Prism 8 software (GraphPad). Differential effects were analyzed by one-way or two-way analysis of variance (ANOVA) followed by appropriate post-hoc tests (Sidak or Dunnett). A *p* value < 0.05 was considered significant (\**p* < 0.05; \*\**p* < 0.01; *** *p* <0.001; \*\*\*\**p* < 0.0001).

## Data availability

The microarray data from this publication have been deposited to the GEO database and assigned the identifier GSE151792 (https://www.ncbi.nlm.nih.gov/geo/query/acc.cgi?acc=GSE151792).

## Acknowledgments

We thank Odile Mondesert and Valerie Lobjois (ITAV, France) for sharing plasmids for chromatibody stable integration and fruitful discussions, and Jacques Dupuy and Fabrice Pierre (Toxalim, France) for sharing the CRISPR plasmid and the HCECs. We thank Hervé Guillou (Toxalim, France) for helpful discussions for transcriptomic data analyzes. This work was supported by the ANR Grant N° ANR-10-CESA-011 to G. Mirey, the Institut National de Recherche pour l’Agriculture, l’Alimentation et l’Environnement (INRAE) and the Toxalim internal program to J. Vignard. B.J. Pons was supported by a Ph.D. fellowship granted by the COLiveTox IDEX Toulouse University.

## Author contributions

B.J. Pons: conceptualization, investigation, methodology, visualization and writing. A. Pettes-Duler, C. Naylies, S. Hashim and S. Tadrist: investigation. F. Taieb: conceptualization, investigation and methodology (*E.col* CDT purification). Y. Lippi: conceptualization, formal analysis, resources, validation, methodology, visualization and writing (transcriptomic analysis). G. Mirey: funding acquisition, project administration and writing. J Vignard: conceptualization, formal analysis, funding acquisition, investigation, methodology, supervision, validation, visualization and writing.

## Supplementary Information

**Fig. S1.**
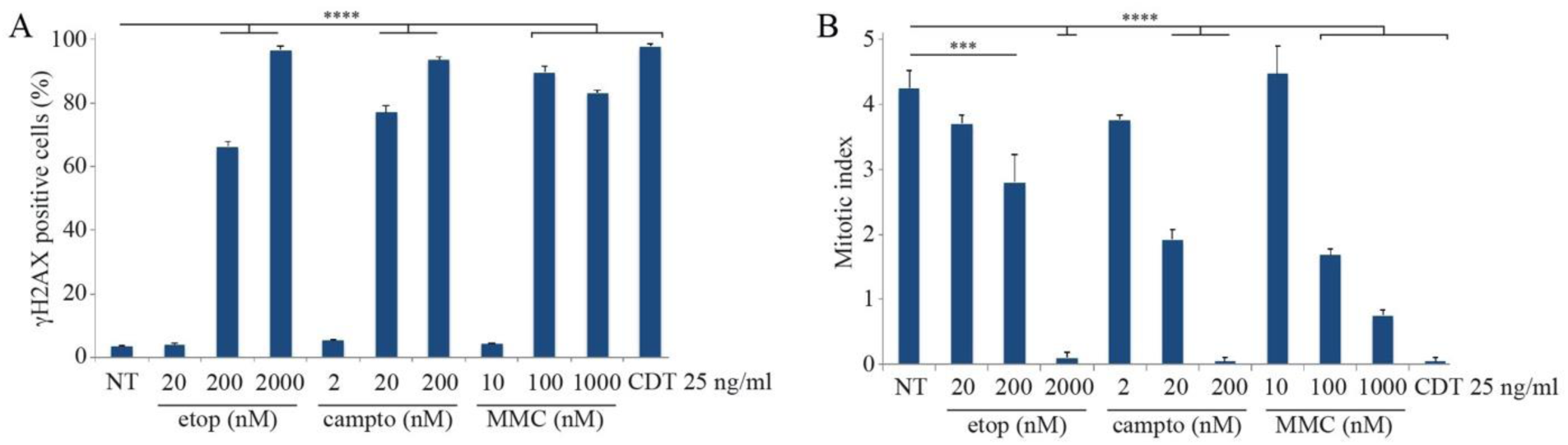
DNA damage and mitotic index in HeLa cells exposed to etop, campto, MMC or CDT 25 ng/ml. (A and B) HeLa cells were exposed for 24 h to etop, campto, MMC or CDT 25 ng/ml and analyzed by immunofluorescence microscopy with an antibody directed against γH2AX. γH2AX positive cells (A) and mitotic index (B) were quantified. Data represent the mean ± SEM (*N*≥3). Statistics were calculated by one-way ANOVA followed by Dunnett’s multiple comparison test.

**Fig. S2.**
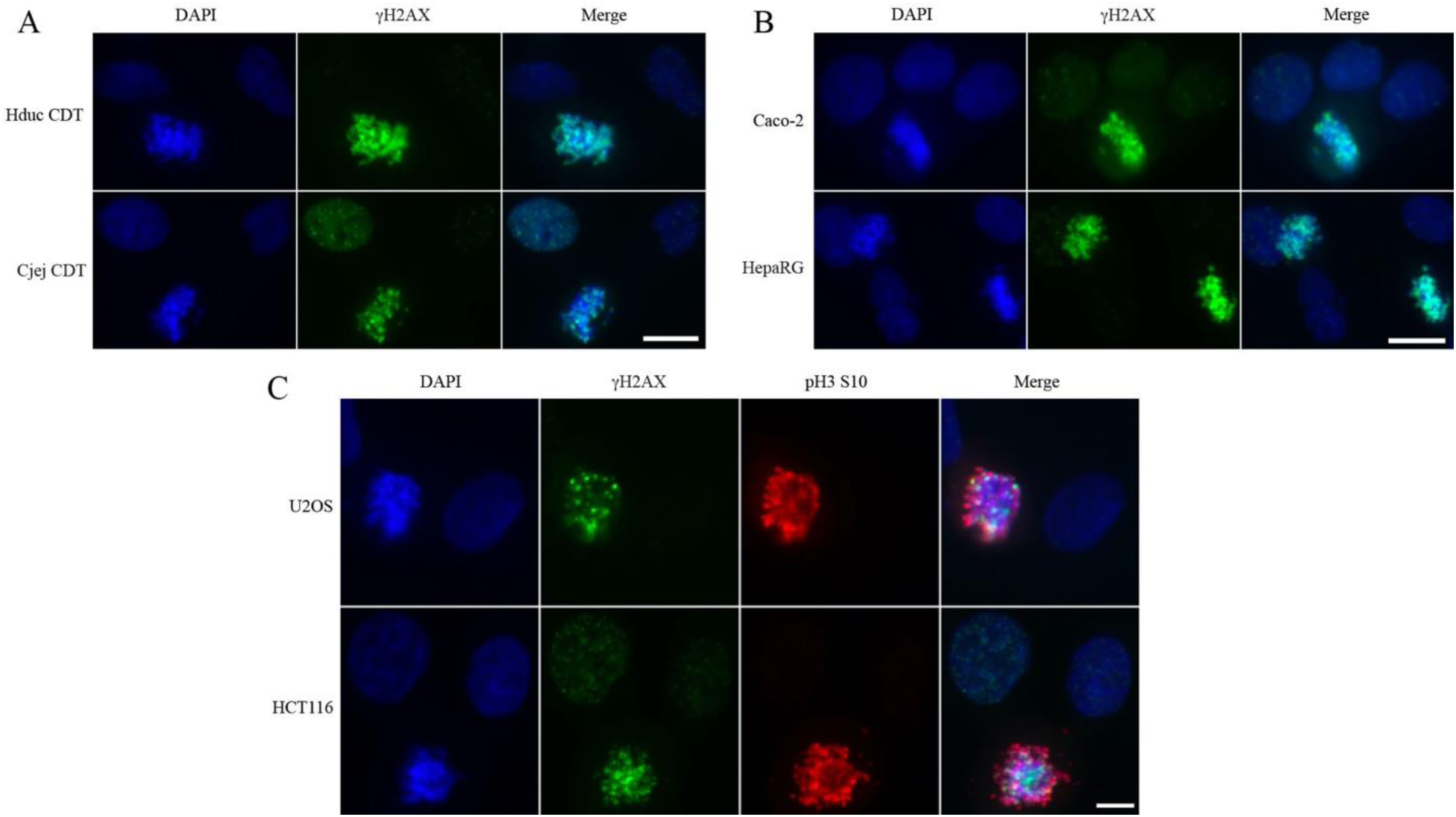
CDT from different origins induce mitotic DNA damage in different cell lines. (**A**) HeLa cells were exposed to Hduc CDT or Cj CDT for 24 h and analyzed by immunofluorescence microscopy with an antibody directed against γH2AX. Scale bar = 20 μm. (**B**) Caco-2 or HepaRG cells were exposed to Ecol CDT for 24 h and analyzed by immunofluorescence microscopy with an antibody directed against γH2AX. Scale bar = 20 μm. (**C**) U2OS or HCT116 cells were exposed to Ecol CDT for 24 h and analyzed by immunofluorescence microscopy with antibodies directed against γH2AX and pH3 S10. Scale bar = 10 μm.

**Fig. S3.**
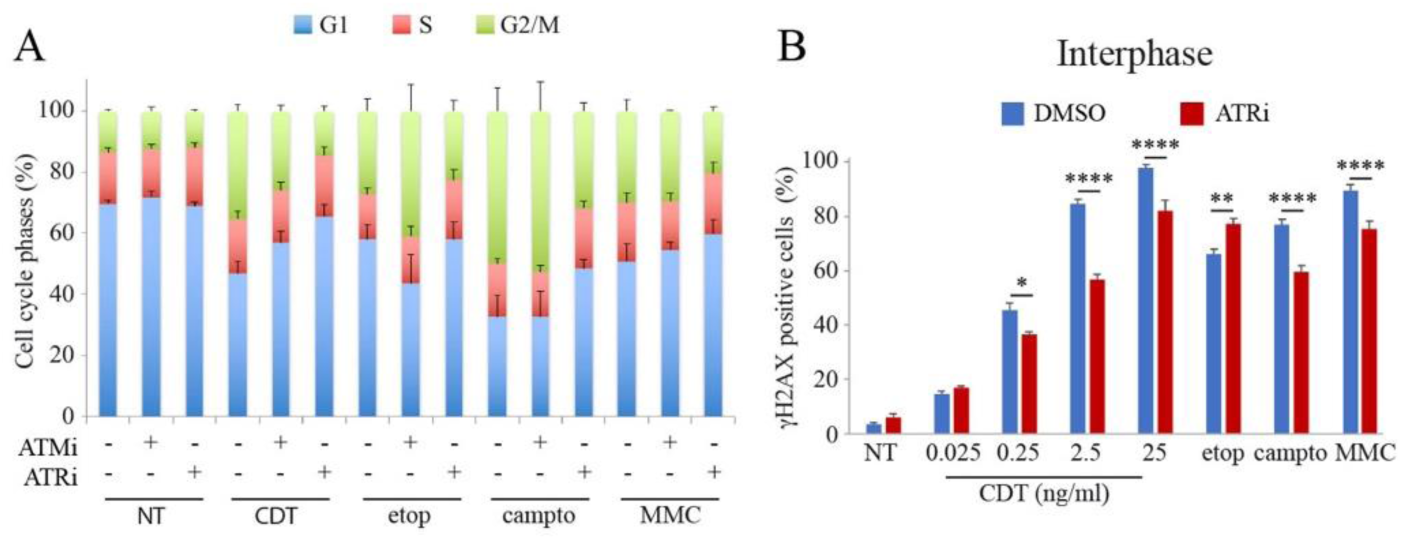
ATR is required for CDT-induced cell cycle arrest and γH2AX increase in interphase. (**A**) HeLa cells were exposed for 24 h to CDT 0.5 ng/ml, etop 200 nM, campto 20 nM or MMC 100 nM with or without ATMi or ATRi and subjected to cell cycle analyzes by flow-cytometry. Data represent the mean ± SEM of at least three independent experiments. (**B**) HeLa cells were exposed for 24 h to CDT, etop 200 nM, campto 20 nM or MMC 100 nM, with or without ATRi, and analyzed by immunofluorescence microscopy with an antibody directed against γH2AX. Data represent the mean ± SEM (*N*≥3). Statistics were calculated by two-way ANOVA followed by Sidak’s multiple comparison test.

**Fig. S4.**
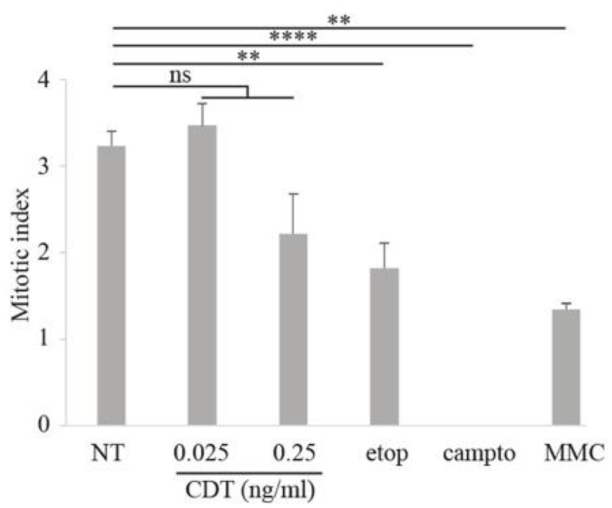
Moderates CDT concentrations do not induce G2 arrest in HCECs. HCECs were exposed for 24 h to CDT, etop 200 nM, campto 20 nM or MMC 100 nM, with or without ATRi, and analyzed by immunofluorescence microscopy with an antibody directed against pH3 S10 for mitotic index quantification. Data represent the mean ± SEM (*N*≥3). Statistics were calculated by two-way ANOVA followed by Sidak’s multiple comparison test.

**Fig. S5.**
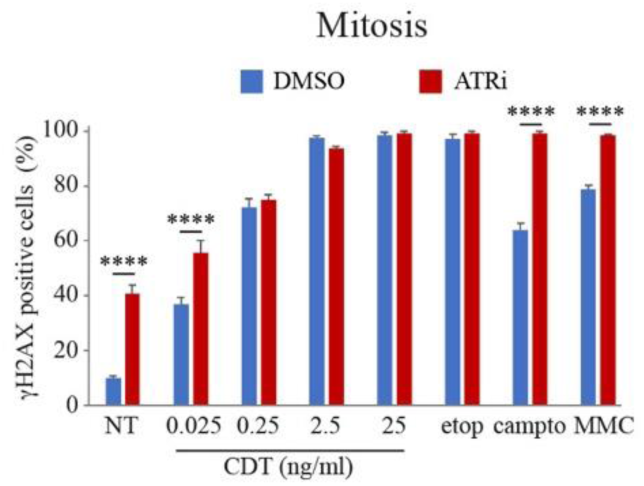
ATR does not prevent the CDT-mediated induction of DNA damage in mitosis. HeLa cells were exposed for 24 h to CDT, etop 200 nM, campto 20 nM or MMC 100 nM, with or without ATRi, and analyzed by immunofluorescence microscopy with antibodies directed against γH2AX and pH3 S10. Only mitotic cells were quantified. Data represent the mean ± SEM (*N*≥3). Statistics were calculated by two-way ANOVA followed by Sidak’s multiple comparison test.

**Fig. S6.**
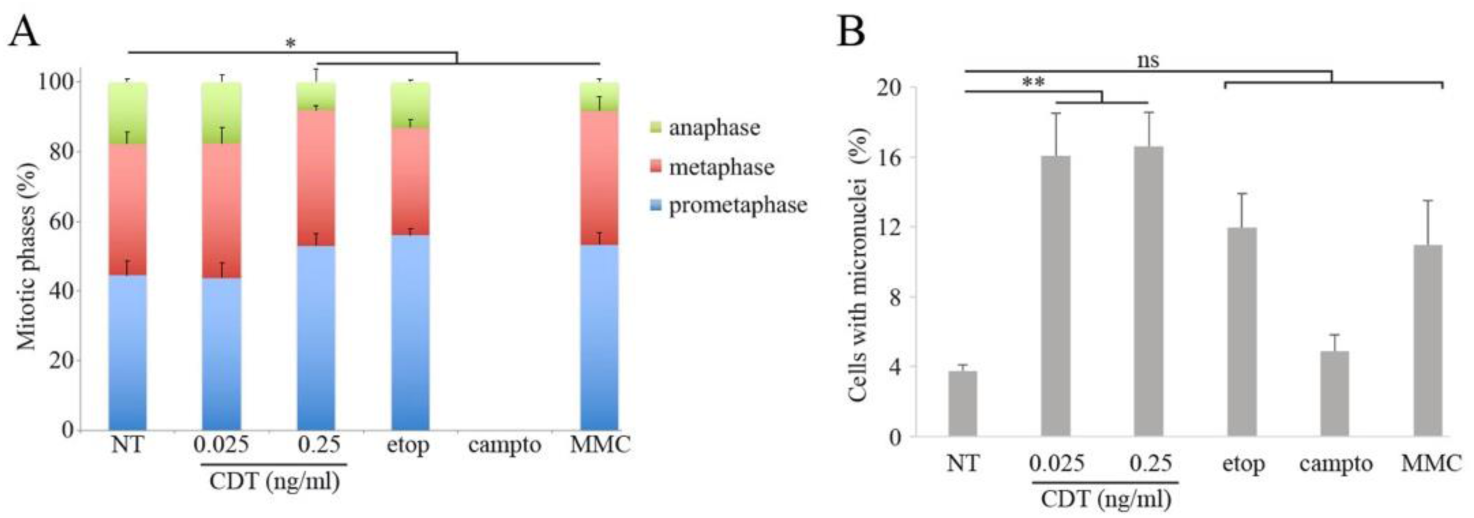
ATR does not prevent the CDT-mediated induction of DNA damage in mitosis. (**A** and **B**) HCECs cells were exposed for 24 h to CDT, etop 200 nM, campto 20 nM or MMC 100 nM and analyzed by immunofluorescence microscopy with an antibody directed against pH3 S10. The proportion of prometaphase, metaphase and anaphase (A), or the frequency of cells with micronuclei (B) were quantified. Data represent the mean ± SEM (*N*≥3). Statistics (only anaphase for A) were calculated by one-way ANOVA followed by Dunnett’s multiple comparison test.

